# LM-GlycoRepo Version 1.0: A novel repository system for mouse tissue glycome mapping data

**DOI:** 10.1101/2025.03.18.644044

**Authors:** Chiaki Nagai-Okatani, Noriaki Fujita, Patcharaporn Boottanun, Miyuki Tanaka, Masaaki Shiota, Daisuke Shinmachi, Kiyohiko Angata, Kiyoko Aoki-Kinoshita, Atsushi Kuno

**Author notes:** **Correspondence:** (C.N.-O.); (A.K.).

## Abstract

Lectin microarray (LMA) is a high-sensitive profiling method of protein glycosylation. The increasing use of this method in many studies has led to a growing demand for a repository system that meets the FAIR data principles (Findable, Accessible, Interoperable, and Reusable). Herein, we present a novel repository system, “LM-GlycoRepo,” for lectin-based multimodal (LM) data, including LMA data, in accordance with the international guideline MIRAGE (Minimum Information Required for a Glycomics Experiment). As a first step in our efforts to provide a general repository for storing various types of LM data, LM-GlycoRepo Version 1.0 is specialized for mouse tissue glycome mapping data obtained using standardized laser microdissection (LMD)-assisted LMA procedures. This system allows users to deposit datasets containing LMD images, LMA data, and high-resolution lectin staining images as LM data. In addition, this repository adopted an “embargo” system that allows users to specify the release date of datasets, allowing compatibility with an article peer review system. Notably, after the release date, the deposited data were visualized using an existing web tool called LM-GlycomeAtlas. LM-GlycoRepo will evolve into a comprehensive tool for lectin-based multimodal data for various biospecimens, including human samples. LM-GlycoRepo is freely available at the GlyCosmos portal (https://lm-glycorepo.glycosmos.org/lm_glycorepo/).

## INTRODUCTION

Protein glycosylation is a major post-translational modification that plays a pivotal role in various biological events^1^. The glycans attached to proteins affect their physiological properties, interactions with other molecules, and molecular recognition, thereby modulating their biological functions. This modification is altered depending on the physiological and pathophysiological state of the host cell, reflecting changes in the glycosylation machinery. Accordingly, analyzing protein glycosylation is important for understanding the biological and pathological roles of glycans and disease-related glycosylation alterations, which offer promising strategies for biomarker and drug development^2,3^. Despite the establishment of many mass spectrometry (MS)-based approaches^4^, analyzing endogenous glycoproteins remains challenging owing to the limitations of sample availability, especially in the case of clinical specimens. In addition, information on whether the target glycans are *N*- and *O*-glycans is also crucial for MS-based glycoproteomics since an appropriate approach is selected depending on the type of glycans^4^. Furthermore, glycosylation is found not only in membrane proteins but also in secretory proteins released from cells, hence, the secreted glycoproteins are not the focus of current single-cell omics technologies^5^. Accordingly, spatial glycomic technology is essential for the comprehensive investigation of endogenous glycoproteins. Although an MS imaging methodology has been established for spatial *N*-glycomics^6^, it cannot yet be applied to spatial *O*-glycomics. Therefore, a highly sensitive analytical methodology is required to obtain spatial information on both *N*- and *O*-glycans, as well as glycoproteins expressed in tissues.

To address this need, lectin microarrays (LMAs) are frequently used to obtain global glycomic profiles of both *N*- and *O*-glycans in biological specimens prior to MS-based analysis that is performed to identify proteins carrying the target glycans and investigate their detailed structures ^7,8^. A major advantage of LMA is its ability to perform highly sensitive and high-throughput differential glycomic analyses using simpler and more rapid procedures than the MS-based approaches. For spatial glycomics, a laser microdissection (LMD)-assisted LMA technique has been standardized to effectively utilize formalin-fixed paraffin-embedded (FFPE) tissues^9^, in which glycans and their carrier proteins are well preserved. Using the standardized procedure^10^, a 0.1 mm^2^ area of 5 μm-thick FFPE tissue sections (approximately 60 cells) is sufficient for differential glycomic profiling with a high-end evanescent-field-activated fluorescence detection scanner^11^. Although it does not provide a detailed structure, this method offers superior sensitivity compared to an MS-based glycomics method combined with LMD-assisted tissue collection for FFPE tissues, which requires 1,000 cells for *N*- and *O*-glycan analysis^12^. Leveraging its sensitivity and reproducibility, the standardized LMD-assisted LMA is used for “tissue glycome mapping,” providing an overview of the quantitative and qualitative distribution of protein glycosylation^9,10,13^. As a feasibility study of the tissue glycome mapping approach, the LMD-assisted LMA method was applied to 14 tissues of normal mice, demonstrating its utility in evaluating and elucidating site- and tissue-specific glycosylation^9,14^. Currently, over 500 data of mouse tissue glycome mapping, along with relevant tissue histological images, are publicly available in the first LMA-based glycomics database, the LM-GlycomeAtlas^14–16^, hosted on the GlyCosmos Portal (https://glycosmos.org/)^17^.

Data sharing based on the FAIR principles (Findable, Accessible, Interoperable and Reusable) is essential for promoting open science^18^. In the field of glycoscience, the MIRAGE (Minimum Information Required for a Glycomics Experiment) initiative was launched in 2011 to develop guidelines for reporting results from various types of glycomics analyses^19^. Guidelines for sample preparation^20^, MS-based glycomics^21^, and glycan microarray^22^ have already been previously published. To support FAIR data sharing in accordance with these guidelines, repository systems for MS-based glycomics (GlycoPOST^23^ and UniCarb-DR^24^) and glycan microarray (within GlyGen^25^) data have been developed. Similarly, the MIRAGE guideline for LMA was released in 2023^26^, as LMA-based glycomics is technically mature and has become a well-accepted method for glycan analysis^7,8^. Although a repository system for protein microarrays is available for LMA data deposition^27^, it does not fully support data deposition or metadata description according to the MIRAGE guideline for LMA.

In this paper, we introduce “LM-GlycoRepo,” a novel repository system for LMA-based glycomics data, which aims to serve as a general repository of lectin-based multimodal (LM) data in accordance with the MIRAGE guidelines. This repository is designed to enable users to deposit and access LM data, including LMD-assisted LMA data and lectin staining images, with relevant metadata sufficient for uses to interpret and reuse the deposited data. As a first step, we developed LM-GlycoRepo Version 1.0, specializing in mouse tissue glycome mapping data obtained via the LMD-assisted LMA method. Users can deposit datasets that include LMD images, LMA data, and high-resolution histological images, such as images of tissue sections stained with multiple lectins on the array. LM-GlycoRepo, as a repository for LMA-based glycomics, was implemented based on the MS-based glycomics repository, GlycoPOST^23^, allowing users to register files and associated metadata in accordance with the MIRAGE guidelines (Figure 1). Another key feature of LM-GlycoRepo Version 1.0 is its ability to visualize deposited data through an updated version of LM-GlycomeAtlas. After the release date specified by the user, the deposited data become becomes visible on the system. In the following, details on the key features of LM-GlycoRepo are provided, with a particular focus on the data submission process and how to use LM-GlycomeAtlas. which is freely available on the GlyCosmos Portal (https://lm-glycorepo.glycosmos.org/lm_glycorepo/).

**Figure 1.**
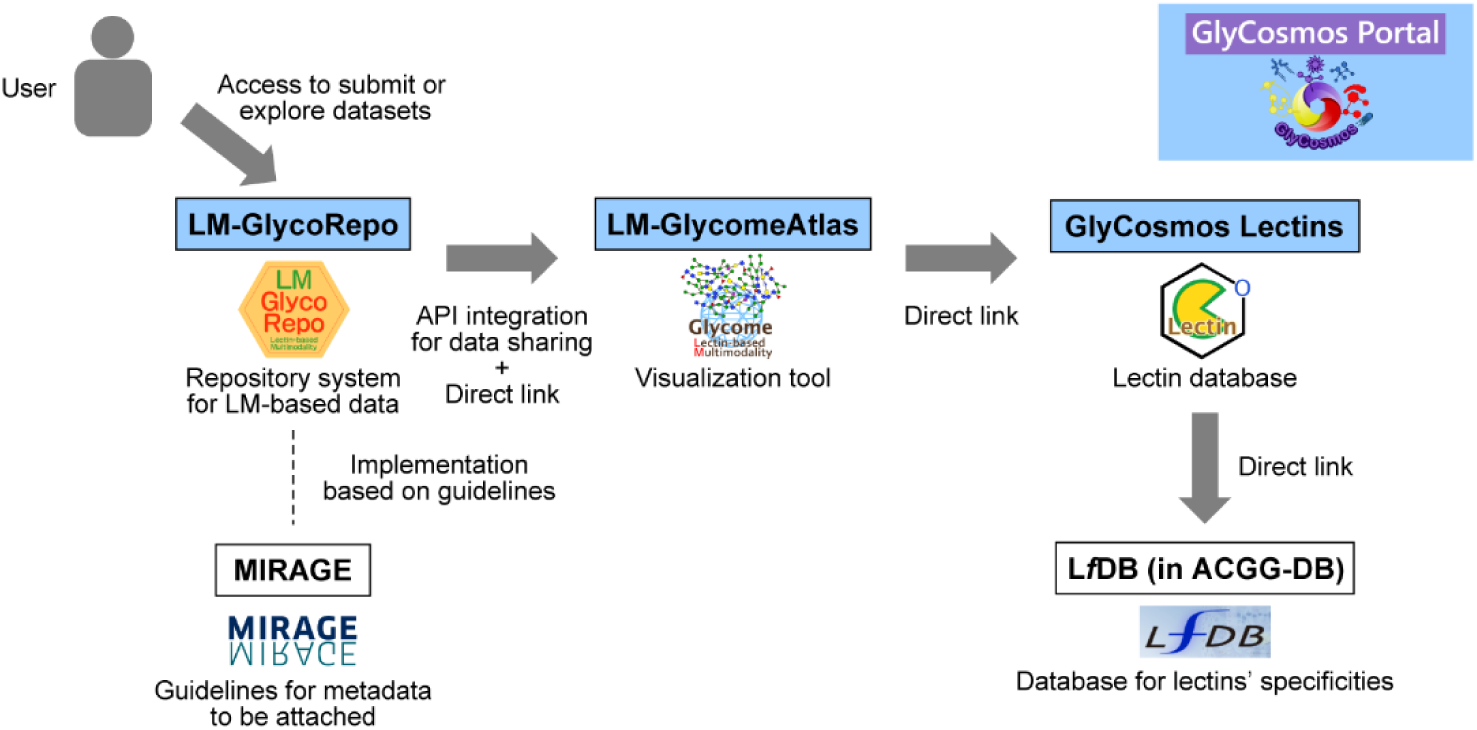
Schematic overview of the relationship between LM-GlycoRepo and its related tools. Users can access LM-GlycoRepo via the GlyCosmos Portal to submit or explore datasets of lectin-based multimodal (LM) data, whose metadata are deposited in accordance with the international guideline, MIRAGE (Minimum Information Required for a Glycomics Experiment). The deposited files and metadata are shared via an application programming interface (API) with LM-GlycomeAtlas, to visualize the data on this web browser-based visualization tool specialized for tissue glycome mapping data. Through LM-GlycomeAtlas, users can access detailed information about the lectins used in the analysis, including their carbohydrate-binding specificities, which are deposited in the L*f*DB, a part of the ACGG-DB, via the lectin database GlyCosmos Lectins.

## METHODS

### Collection of mouse tissue glycome mapping datasets

Four published datasets available in LM-GlycomeAtlas Version 2.1 were used to construct LM-GlycoRepo and are referred to as Reference 1–4, respectively. The datasets of Reference 1^14^ and 2^9^ were obtained from FFPE sections of nine (pancreas, heart, lung, thymus, gallbladder, stomach, small intestine, colon, and skin) and five (brain, liver, spleen, kidney, and testis) tissues, respectively, from 8-week-old normal male C57BL/6J mice. These glycomic profiling data from 14 tissues were collected to provide an overview of site- and tissue-specific glycosylations. The dataset of Reference 3^28^ was obtained from the cardiac tissues of 24-week-old dilated cardiomyopathy (DCM) model and age-matched normal mice, whose glycomic profiles have been used to discover DCM-associated aberrant glycosylations. Reference 4^16^ includes a dataset obtained from the cardiac tissues of C57BL/6N female mice at three different ages (2, 12–14, and 23–25 months), whose glycomic profiles have been compared to elucidate age-related glycosylation changes. Notably, for Reference 3, tissue glycome mapping data were accompanied by high-resolution images of coronal cardiac sections fluorescently stained with nine lectins (i.e., PSA, LCA, AOL, AAL, MAL-I, LEL, ACA, MAH, and WFA) selected on the basis of tissue glycome mapping results. The dataset also includes tissue staining images for hematoxylin and eosin (HE) and collagen staining dyes (Masson’s trichrome and picrosirius red staining), allowing for morphological observation and fibrosis detection, respectively.

For all four datasets, tissue glycome mapping data were obtained using a standardized LMD-LMA procedure, as previously described^10^. Briefly, tissue fragments were collected from a small area (0.13–0.91 mm2) of hematoxylin-stained FFPE tissue sections (5 μm-thick) using an LMD system (LMD6500; Leica Microsystems, Wetzlar, Germany) and subjected to protein extraction with 0.5% Nonidet P-40 as a detergent. The resulting protein extracts were fluorescently labeled with a Cy3-dye and then subjected to LMA using a commercial array chip (LecChip Ver. 1.0; GlycoTechnica, Yokohama, Japan) equipped with 45 lectins (Table S1; where the abbreviations of lectins are provided) and an evanescent field fluorescence scanner (GlycoStation Reader 1200; GlycoTechnica). Note that LecChip Ver.1.0 has been discontinued, but its compatible product, LecChip 45-uni, is currently supplied by Precision System Science (Matsudo, Japan). The resulting glycomic profiles are presented as the signal intensities of the 45 lectins, normalized to the average intensity for all lectins.

### System implementation

The LM-GlycoRepo web application includes a simple user interface for data registration and an application programming interface (API) program that stores information and images in a database or file. The user interface was developed using JSP and Java based on that of GlycoPOST^23^. The API program was developed using the Apache Spark framework (https://spark.apache.org/) with PostgreSQL (https://www.postgresql.org/) for database management, to provide a representational state-transfer (REST) API. The API was used for exclusively for data sharing with LM-GlycomeAtlas. Accordingly, the data storage and user interface programs are independent of each other, allowing for easier development. In addition, LM-GlycomeAtlas, a highly usable web application, can display data updated by LM-GlycoRepo without knowing the state of the data required for the display.

### MIRAGE guidelines

To enable data deposition in compliance with the FAIR principles, the information that should be registered in LM-GlycoRepo Version 1.0 was defined according to the MIRAGE guidelines (https://www.beilstein-institut.de/en/projects/mirage/guidelines/) for sample preparation (version 1.0; doi:10.3762/mirage.1) and LMA (version 1.0; doi:10.3762/mirage.8).

## RESULTS AND DISCUSSION

### Compliance with the MIRAGE guidelines

As mentioned in the Introduction, we aimed to design LM-GlycoRepo for the deposition of metadata in accordance with the MIRAGE guidelines. These metadata are important for users to assess the adequacy of analytical methods and to interpret the biological and pathological significance. Therefore, the items to be registered at each step were determined based on the guidelines for LMA, which require the description of the sample source and preparation, lectin library, array fabrication, data acquisition protocol including the detector used, and data processing and presentation. The items to be included in the sample description were also decided by referring to the guidelines for sample preparation. Accordingly, in the submission system, these items specified in the guidelines were set as mandatory items so that registration cannot be completed unless they are entered, ensuring the attachment of the metadata defined in the guidelines.

### How to submit data on LM-GlycoRepo

Data deposition on LM-GlycoRepo involves two main steps: “Create Template” and “Register Project” (Figure 2). In the first step, users access the “Create Template” section to set up templates detailing the experimental procedures, including tissue section preparation, LMD-assisted sample collection, and subsequent LMA. These templates are then used in the “Register Project” section to simplify the entry of metadata associated with the uploaded data files. Once created, the templates can be reused with modification (if necessary) for future projects with similar experimental procedures.

**Figure 2.**
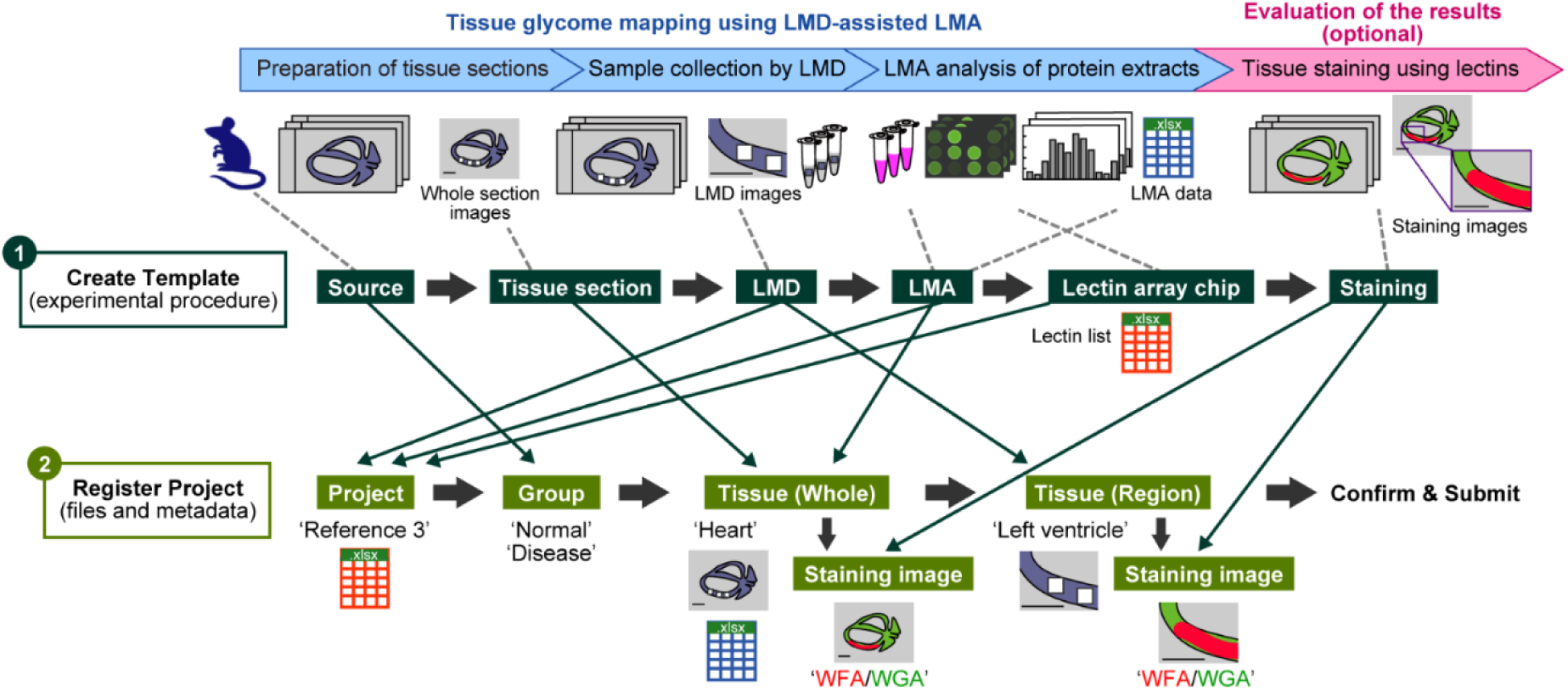
Schematic overview of the data submission process on LM-GlycoRepo. This system allows users to upload tissue glycome mapping data obtained via LMD-assisted LMA, along with optional tissue staining data. The process starts in the “Create Template” section, where users generate six types of templates covering source materials and experimental procedures. Next, in the “Register Project” section, users upload their files and metadata using the previously created templates. The figure illustrates a dataset from Reference 3, which includes two groups—normal and diseased mice—with LMD-assisted LMA data from the left ventricle and tissue images of heart sections stained with lectins such as WFA and WGA.

#### 1. Metadata preparation in the “Create Template” section

The “Create Template” section contains six subsections: (1) Source, (2) Tissue sections, (3) LMD, (4) LMA, (5) Lectin array chip, and (6) Staining (Figure 2). Details on each of these subsections are provided below.

**1) Source.** This subsection describes the creation of templates containing information on biological sources including species, strain, age, and sex. In version 1.0, only the mouse data can be registered.
**2) Tissue section.** This subsection describes all aspects of tissue section preparation, including the section type (e.g., FFPE), fixation method (e.g., 4% paraformaldehyde), section thickness, slide glass type (e.g., PEN), and LMD staining method (e.g., hematoxylin). Detailed staining methods follow the format of the Experimental Procedures section of the research paper.
**3) LMD.** This subsection describes the creation of templates describing laser microdissection (LMD) procedures, including the supplier and instrument name of the LMD system, and the type of observation (e.g., bright field). The detailed LMD procedures are also described in the Experimental Procedures section of a research paper.
**4) LMA.** This subsection covers all aspects of the lectin microarray (LMA) procedures, except for the lectin array chip, which is described in the following subsection. Information such as the supplier and instrument name of the LMA system and the image analysis software for the LMA scans should be included. The detailed LMA procedures, including sample preparation, data acquisition, processing, and statistical analysis, are also described as in the Experimental Procedures section of a research paper.
**5) Lectin array chip.** This subsection describes detailed information regarding lectin array chips.

The information entered was used directly for project registration without modification. Because the items to be registered varies depending on whether the chips are custom-made or commercial, users first select the chip type.

For custom-made chips, details should be provided according to the MIRAGE guidelines for LMA, covering sections such as “immobilization surface” and “array production,” which includes the details of quality control.

For commercial chips, the inclusion of array fabrication details is optional (except for the number of lectin replicates, e.g., three spots). Instead, product identification details, including name, supplier, catalog number, and website URL, should be provided, allowing for users to access to the information about the chip including the quality control that has been performed by the supplier.

A list of lectins used in the array panel should also be provided and formatted as a template (.xlsx), including lectin names and their identifiers (i.e., UniProt ID or GlyCosmos Lectin No.) (Table S1). It is recommended to use lectin names provided by the supplier be used for consistency, especially for commercial chips. The identifiers are used for automatically making a direct link to the lectin’s detail page of the GlyCosmos Lectins (https://glycosmos.org/lectins) on the main page of the LM-GlycomeAtlas. For custom-made chips, additional information is required to comply with the MIRAGE guidelines, including the vender, type of source (i.e., natural or recombinant), modification, and a lot of lectins used for the fabrication of LMA, allowing for users to check the details of the array used for analysis.

**1) 6) Staining.** This subsection describes all aspects of the tissue staining procedures that may accompany tissue glycome mapping experiments. This can include conventional methods, such as hematoxylin and eosin (HE) staining for morphological observation or histochemical staining using lectins. Information such as the method name (e.g., HE) or lectin name (e.g., AAL), along with the supplier and product details of the microscopy system used for imaging, should be included. Detailed staining protocols and microscopic methods are also described as in the Experimental Procedures section of a research paper.

#### 2. Data registration in the “Register Project” section

This section comprises four subsections: (1) Project, (2) Group, (3) Tissue (Whole), and (4) Tissue (Region) (Figure 2). Steps 3 and 4 include an optional module for registering the staining images. After providing all required information, registrants can review the uploaded files and metadata before finalizing the submission. As an “embargo” system, when finalizing the submission, registrants can determine when the deposited data will be released to the public. Details on each subsection of the submission process are provided below.

**1) Project.** In this subsection, users define a project name that serves as the primary identifier in the data list displayed on the LM-GlycoRepo homepage. Users also register the details about the LMD and LMA procedures and the lectin array chip used for the project, using templates created in the “Create Template” section. Since this system allows only one setting for these details, the data set obtained using a different experimental procedure and lectin array chip should be registered as a different project.
**2) Group.** In this subsection, users define the sample groups to be compared (e.g., “Normal” and “Disease”). Users can select a preset “Source” template to autofill in the biological source information, such as strain, age, and sex. Additionally, the number of individuals used for the analysis should be provided.
**3) Tissue (Whole).** This subsection is for registering information about tissue sections analyzed in a project, allowing users to autofill details using the “Tissue Section” template. Users must upload a whole-section image file (in .png format), in which the regions used for tissue collection are labeled. An LMA data file (in .xlsx format) should also be uploaded for each tissue, ensuring that all LMA data registered within a single project are comparable (i.e., array format and chip lot should be identical). As an option, users can register staining images for whole tissue sections and the associated metadata by using a “Staining” template.
**4) Tissue (Region).** In this subsection, users upload a composite image file that summarizes multiple LMD images for a single tissue region (e.g., the left ventricle of the heart). This section also allows users to register detailed information and files for magnified tissue staining images for a single region, similar to the “Tissue (Whole)” section.

### Visualization of the deposited data on LM-GlycoRepo

As the “embargo” system, after the registrant-specified release date, the submitted files and metadata are made visible on a data list of the LM-GlycoRepo homepage. Other users can access and download the deposited resources as they were submitted from the “Detail page”, where all deposited files and metadata are available, allowing for users to review the details of the source and experimental procedures and to re-analyze offline.

Upon public release, LM-GlycoRepo Version 1.0 contains four projects linked to previously published articles, denoted as References 1–4. These tissue glycome mapping datasets were obtained from 14 tissues of normal and diseased mice using LMD-assisted LMA with 45 lectins (Table S1) and were available in an earlier version (Version 2.1) of the LM-GlycomeAtlas. The glycomic profiles obtained from normal C57BL/6J male mice in Reference 1^14^ and Reference 2^9^ are useful for an overview of site- and tissue-specific tissue glycosylations. For example, a previous study employed these data to evaluate MS-based glycoproteomics results for elucidation of inter-tissue glycan heterogeneities and to obtain additional spatial information of target glycans within a tissue^29^. Reference 3^28^ contains a data set of cardiac tissues from the DCM model and age-matched normal mice used to elucidate glycosylation changes associated with cardiac fibrosis in failing hearts. Notably, this study identified WFA as a lectin suitable for the detection of cardiac fibrogenesis, which can be verified by comparing the high-resolution images of collagen staining (i.e., MT and PSR) and WFA staining using the viewer provided with LM-GlycomeAtlas. Reference 4^16^ is a data set of cardiac tissues obtained from normal C57BL6/N female mice at three different ages, providing insight into region-specific glycosylation and its age-related changes.

As mentioned above, a key feature of LM-GlycoRepo is the visualization of deposited data on LM-GlycomeAtlas by API data sharing (Figure 2). To this end, each entry in the data list of LM-GlycoRepo contains a direct link to the LM-GlycomeAtlas, allowing users to visually explore the data using the web-based tools provided on the GlyCosmos Portal (Figure 1), as described below.

### Visualization of the deposited data on LM-GlycomeAtlas

To enhance the visualization of tissue glycome mapping data deposited in LM-GlycoRepo Version 1.0, the user interface of LM-GlycomeAtlas Version 2.0^15^ was updated to Version 2.2 with several improvements. One major enhancement is the display of the metadata registered on LM-GlycoRepo via API integration. The data integration is automatic and therefore this system does not provide a way for users to access the API. On the main page of the LM-GlycomeAtlas Version 2.2, a tissue chart shows project-specific information, including the project name, group name, number of individuals analyzed, and number of deposited histological images (Figure 3A). After selecting the tissue of interest, a whole-tissue section image uploaded by the user is displayed, allowing users to review the histology of the target tissue (Figure 3B). Simultaneously, a graph appears below the chart, showing the LMA data for all samples from the selected tissue (Figure 3C). To access additional details, users can interact with the functional buttons below the whole-tissue section image (Figure 3B). When a region of interest is selected using the “Region” button, the graph updates to display LMA data for the selected region only (Figure 3C). Additionally, an LMD image of the selected region uploaded by the user is displayed in a new window (Figure 4D), allowing users to examine the morphology of the tissues used for LMA analysis. If the project includes histological images, the “Staining” button will become available under the “Region” button (Figure 3B), enabling users to view multiple selected images. These images are displayed in separate windows along with metadata, including the staining methods (e.g., lectin staining), observation methods (e.g., fluorescence), and the staining dye used (e.g., Blue: Hoechst33342) (Figure 3E). This visualization system supports high-resolution images with scalable zoom, enabling users to examine stained cells and tissues in detail.

**Figure 3.**
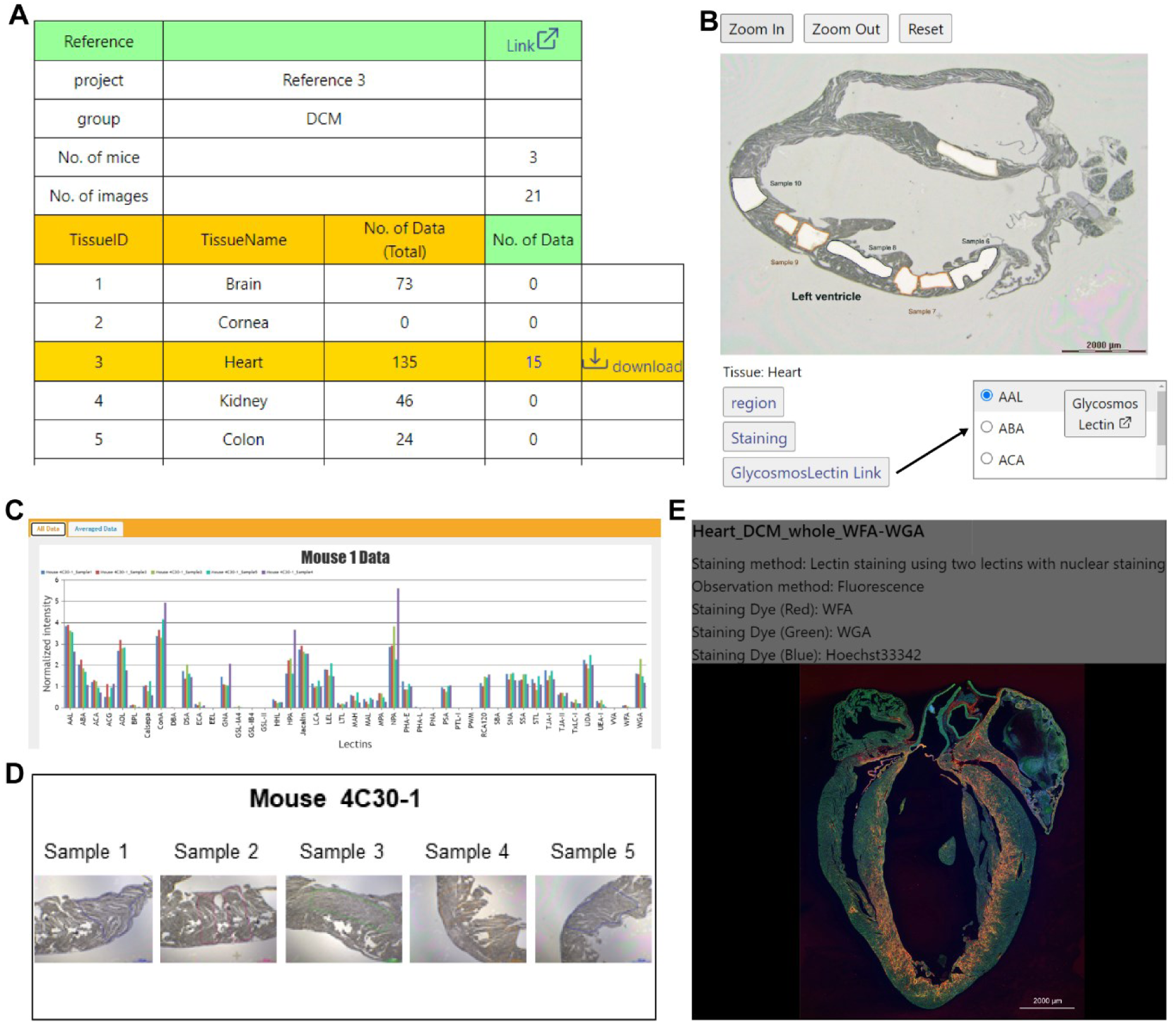
Representative images of the LM-GlycomeAtlas Version 2.2 user interface. The main page shows the tissue list and project information (A), tissue section image and functional buttons (B), and a graph showing LMA data (C). Users can visualize an LMD image (D) and tissue staining image (E) in separate windows.

Another key feature of LM-GlycomeAtlas Version 2.2 is the direct link to the lectin database “GlyCosmos Lectins” (Figure 1). The main page contains a “GlyCosmos Lectin Link” button (Figure 3B) that allows users to select a lectin of interest from an array panel. The GlyCosmos Lectins, a resource available on the GlyCosmos Portal, offers a comprehensive list of glycan-binding proteins, including lectins. Each entry provides basic information, such as the source organism and monosaccharide specificity, as well as a direct link to databases containing glycan interaction data, where available. For example, the Lectin Frontier DataBase (LfDB) within the framework of the Asian Community of Glycoscience and Glycotechnology Database (ACGG-DB) provides interaction data between lectins and pyridylaminated glycans using frontal affinity chromatography with fluorescence detection^30^ (Figure 1). Other databases, such as MCAW-DB^31^ and CarboGlove^32^, which offer glycan-binding specificities obtained using glycan microarrays, are also accessible via GlyCosmos Lectins. This streamlined access to the carbohydrate specificity data of lectins used in the LMA analysis aims to help users interpret the tissue glycome mapping data available on the LM-GlycomeAtlas.

### Limitations and Perspectives

Our goal is to construct a repository system that allows the deposit of detailed metadata entries in a format suitable for an experimental method used for analysis. This point is essential to achieve both the detailed metadata description in accordance with the MIRAGE guidelines and the easy exploration of the deposited data by other users. To achieve these goals, we have adopted a strategy of starting with LMD-assisted LMA data (associated with lectin staining images) of mouse tissues as a dataset of LM data with metadata compliant with the guidelines. Owing to this bottom-up strategy, LM-GlycoRepo Version 1.0 has several limitations.

A major limitation is that this repository only covers the mouse species to prioritize the visualization of deposited data on LM-GlycomeAtlas, which currently specializes in mouse data. The LM-GlycoRepo system itself may cover other species only with slight modification. In addition, the LM-GlycomeAtlas system, which is based on the GlycomeAtlas system, is highly extensible, as GlycomeAtlas covers human and zebrafish data in addition to mouse data^33^ (Figure 1). Leveraging these features, both LM-GlycoRepo and LM-GlycomeAtlas have the potential to expand the range of data types deposited and visualized, respectively.

Another major limitation is that the data that can be deposited is limited to glycomic profiles obtained from FFPE tissue sections using LMD-assisted LMA. To serve as a general repository for LM data, LM-GlycoRepo should be improved in the future to enable the deposit of data obtained by other methods. For example, macro-dissection is an alternative method for obtaining tissue samples that should be covered. In addition, spatial glycomics using MS-based methods such as imaging mass cytometry and MS imaging are useful for high-resolution and high field-of-view analyses, respectively^34^. Accordingly, the integration with existing initiatives to build an atlas specialized for MS imaging data^35^ can offer further interactions.

## CONCLUSIONS

As the fourth repository system on the GlyCosmos Portal, LM-GlycoRepo Version 1.0 represents the first repository to comply with the MIRAGE guidelines for LMA-based glycomics data, facilitating data sharing under the FAIR principles. These LMA-based spatial glycomics data are valuable for complementing and enhancing MS-based approaches for glycomics^36^ and glycoproteomics^37,38^ of mouse tissues, especially to gain insights into inter-tissue glycan heterogeneities^29^. The GlyCosmos Portal makes glycoconjugates available as a common “language” by assigning specific identifiers to them in a standardized repository called GlyComb^39^. By leveraging this framework, LM-GlycoRepo, LM-GlycomeAtlas, and their relevant tools in the GlyCosmos Portal (Figure 1) can interactively provide lectin interaction-based glycan analysis data.

## ASSOCIATED CONTENT

### Supporting Information

The following files are available free of charge.

**Table S1.** Abbreviations and carbohydrate specificities of 45 lectins on LecChip Ver. 1.0 (PDF)

### Data Availability Statement

All data described here are freely available on LM-GlycoRepo (https://lm-glycorepo.glycosmos.org/lm_glycorepo/).

## AUTHOR INFORMATION

### Corresponding Authors

(C.N.-O.); (A.K.)

### Notes

The authors declare no competing financial interest.

## ACKNOWLEDGMENTS

We are grateful to Dr. Shujiro Okuda of Niigata University for the provision of the GlycoPOST source code. This work was supported by the Database Integration and Coordination Program (DICP) sponsored by the Japan Science and Technology Agency (JST) and the National Bioscience Database Center (NBDC) (grant number: JPMJND2204). Authors from GaLSIC have been partly supported by J-GlycoNet, one of the official Joint Usage/Research Centers of the Ministry of Education, Culture, Sports, Science and Technology (MEXT) in Japan.

## For TOC Only

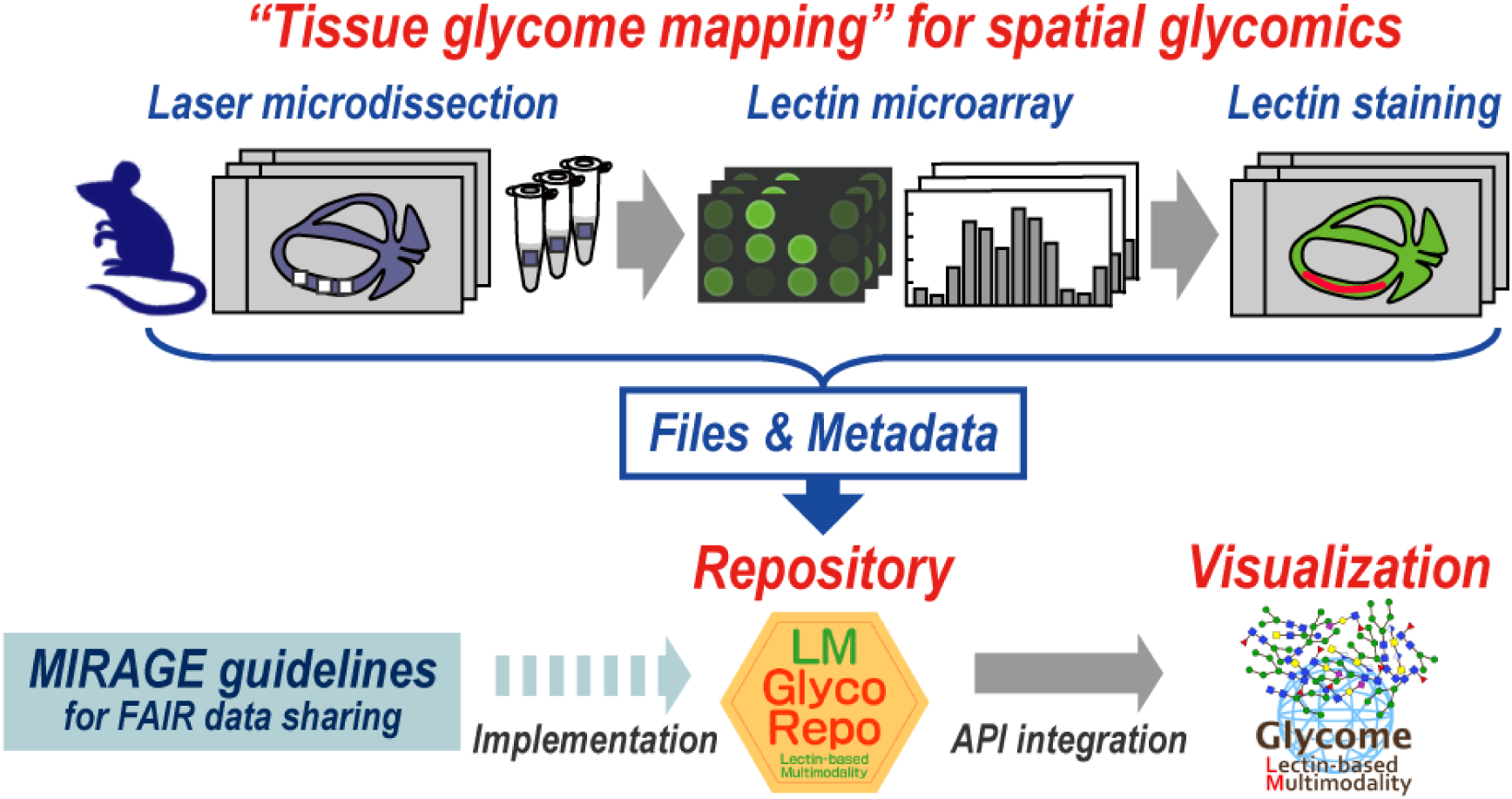

