## Supporting Information for "LM-GlycoRepo Version 1.0: A novel repository system for mouse tissue glycome mapping data"

### LM-GlycoRepo Version 1.0: A novel repository system for lectin microarray-based mouse tissue glycome mapping data

#### Table of Contents

**Table S1.** Abbreviations and carbohydrate specificities of 45 lectins on LecChip Ver. 1.0.

**Table S1. Abbreviations and carbohydrate specificities of 45 lectins on LecChip Ver. 1.0.***

| | Lectin | Origin | Binding specificity** | Uniprot ID | GlyCosmos Lectin No. |
|---|---|---|---|---|---|
| 1 | AAL | *Aleuria aurantia* | Terminal α-Fuc, Sia-$Le^x$ and $Le^x$ | P18891 | GL_000407 |
| 2 | ABA | *Agaricus bisporus* | Galβ1-3GalNAcα-Thr/Ser (T) and sialyl-T | Q00022 | GL_000025 |
| 3 | ACA | *Amaranthus caudatus* | Galβ1-3GalNAcα-Thr/Ser (T) | Q71QF2 | GL_002234 |
| 4 | ACG | *Agrocybe cylindracea* | Siaα2-3Galβ1-4GlcNAc | A0A2D0TCI3 | GL_001912 |
| 5 | AOL | *Aspergillus oryzae* | Terminal α-Fuc, Sia-$Le^x$ and $Le^x$ | Q2UNX8 | GL_002011 |
| 6 | BPL | *Bauhinia purpurea alba* | Galβ1-3GalNAc and $NA_3$, $NA_4$ | P16030 | GL_000686 |
| 7 | Calsepa | *Calystega sepiem* | Man and Maltose | P93114 | GL_000600 |
| 8 | ConA | *Canavalia ensiformis* | α-Man (inhibited by presence of bisecting GlcNAc) | P02866 | GL_002303 |
| 9 | DBA | *Dolichos biflorus* | GalNAcα-Thr/Ser (Tn) and GalNAcα1-3GalNAc | P05045 | GL_002320 |
| 10 | DSA | *Datura stramonium* | (GlcNAc)n, polyLacNAc and LacNAc ($NA_3$, $NA_4$) | A0A089ZWN7 | GL_001968 |
| 11 | ECA | *Erythrina cristagalli* | Lac/LacNAc | P83410 | GL_000692 |
| 12 | EEL | *Euonymus europaeus* | Galα1-3[Fucα1-2Gal] > Galα1-3Gal | B3SV75 | GL_002417 |
| 13 | GNA | *Galanthus nivalis* | Non-substituted α1-6Man | P30617 | GL_002361 |
| 14 | GSL-I A4 | *Griffonia simplicifolia* | α-GalNAc and GalNAcα-Thr/Ser (Tn) | Q8W1R7 | GL_001974 |
| 15 | GSL-l B4 | *Griffonia simplicifolia* | α-Gal | Q8W1R6 | GL_001959 |
| 16 | GSL-II | *Griffonia simplicifolia* | Agalactosylated *N*-glycan | Q41263 | GL_001972 |
| 17 | HHL | *Hippeastrum hybrid* | Non-substituted α1-6Man | - | GL_002430 |
| 18 | HPA | *Helix pomatia* | Terminal GalNAc | Q2F1K8 | GL_001939 |
| 19 | Jacalin | *Artocarpus integrifolia* | Galβ1-3GalNAcα-Thr/Ser (T) and GalNAcα-Thr/Ser (Tn) | P18670 | GL_000029 |
| 20 | LCA | *Lens culinaris* | Fucα1-6GlcNAc, α-Man and α-Glc | P02870 | GL_001682 |
| 21 | LEL | *Lycopersicon esculentum* | (GlcNAc)n and polyLacNAc | G9M5T0 | GL_002419 |
| 22 | LTL | *Lotus tetragonolobus* | Fucα1-3GlcNAc, Sia-$Le^x$ and $Le^x$ | P19664 | GL_000058 |
| 23 | MAH | *Maackia amurensis* | Siaα2-3Galβ1-3[Siaα2-6GalNAc]α-R | Q7M1M0 | GL_001973 |
| 24 | MAL I | *Maackia amurensis* | Siaα2-3Gal | P0DKL3 | GL_001099 |
| 25 | MPA | *Maclura pomifera* | Galβ1-3GalNAcα-Thr/Ser (T) and GalNAcα-Thr/Ser (Tn) | P18674 | GL_000030 |
| 26 | NPA | *Narcissus pseudonarcissus* | Non-substituted α1-6Man | C0HM45 | GL_002418 |
| 27 | PHA-E | *Phaseolus vulgaris* | $NA_2$ and bisecting GlcNAc | P05088 | GL_002322 |
| 28 | PHA-L | *Phaseolus vulgaris* | Tri- and tetra-antennary complex-type *N*-glycan | P05087 | GL_002321 |
| 29 | PNA | *Arachis hypogaea* | Galβ1-3GalNAcα-Thr/Ser (T) | P02872 | GL_000456 |
| 30 | PSA | *Pisum sativum* | Fucα1-6GlcNAc and α-Man | P02867 | GL_002304 |
| 31 | PTL-I | *Psophocarpus tetragonolobus* | α-GalNAc and Gal | - | GL_002433 |
| 32 | PWM | *Phytolacca americana* | (GlcNAc)n and polyLacNAc | P83790 | GL_000736 |
| 33 | RCA120 | *Ricinus communis* | Lac/LacNAc | P02879 | GL_001077 |
| 34 | SBA | *Glycine max* | Terminal GalNAc (especially GalNAcα1-3Gal) | P05046 | GL_000685 |
| 35 | SNA | *Sambucus nigra* | Siaα2-6Gal/GalNAc | Q41358 | GL_001074 |
| 36 | SSA | *Sambucus sieboldiana* | Siaα2-6Gal/GalNAc | Q58A86 | GL_002423 |
| 37 | STL | *Solanum tuberosum* | (GlcNAc)n and polyLacNAc | Q9S8M0 | GL_000308 |
| 38 | TJA-I | *Trichosanthes japonica* | Siaα2-3Galβ1-4GlcNAcβ-R | - | GL_002429 |
| 39 | TJA-II | *Trichosanthes japonica* | Fucα1-2Gal, β-GalNAc > $NA_3$, $NA_4$ | - | GL_002432 |
| 40 | TxLCI | *Tulipa gesneriana* | Man3, bi- and tri-antennary complex-type *N*-glycan, GalNAc | - | GL_002431 |
| 41 | UDA | *Urtica dioica* | (GlcNAc)n and polyLacNAc | P11218 | GL_000759 |
| 42 | UEA I | *Ulex europaeus* | Fucα1-2LacNAc | P22972 | GL_000060 |
| 43 | VVA | *Vicia villosa* | α-, β-linked terminal GalNAc and GalNAcα-Thr/Ser (Tn) | P56625 | GL_000721 |
| 44 | WFA | *Wisteria floribunda* | Terminal GalNAc (e.g., GalNAc β1-4GlcNAc) | A0A218PFP3 | GL_001969 |
| 45 | WGA | *Triticum vulgaris* | (GlcNAc)n and multivalent Sia | P10969 | GL_000044 |

*LecChip Ver. 1.0 is a commercialized lectin array chip provided by GlycoTechnica Ltd.

**Binding specificities are based on Lectin Frontier Database (LfDB; https://acgg.asia/lfdb2).
